# Australian vertebrate hosts of Japanese encephalitis virus; a review of the evidence

**DOI:** 10.1101/2024.04.23.590833

**Authors:** Kevin T. Moore, Madelyn J. Mangan, Belinda Linnegar, Tejas S. Athni, Hamish I. McCallum, Brendan J. Trewin, Eloise Skinner

**Affiliations:** Centre for Planetary Health and Food Security, Griffith University, Gold Coast, QLD 4222, Australia; Harvard Medical School, Boston, MA 02115, USA; CSIRO, Health and Biosecurity, Brisbane, QLD 4102, Australia; Department of Biology, Stanford University, Stanford, CA 94305, USA

**Keywords:** Host competence, Maintenance host, Spillover host, Transmission cycle, Experimental infection, Viraemia

## Abstract

Japanese Encephalitis Virus (JEV) transmission in temperate Australia has underscored a critical need to characterise transmission pathways and identify probable hosts of infection within the country. This systematic review consolidates existing research on the vertebrate hosts of JEV that are known to exist in Australia. Specifically, we aim to identify probable species for JEV transmission, their potential role as either a spillover or maintenance host and identify critical knowledge gaps. Data were extracted from studies involving experimental infection, seroprevalence, and virus isolation and were available for 22 vertebrate species known to reside in Australia. A host competence score was calculated to assess the potential for a given species to infect JEV vectors and to quantity their possible role in JEV transmission. Based on the host competence score and ecology of each species, we find ardeid birds, feral pigs, and flying foxes have potential as maintenance hosts for JEV in the Australian context. We also note that brushtail possums and domestic pigs have potential as spillover hosts under certain outbreak conditions. However, evidence to confirm these roles in localized transmission or outbreaks is sparse, emphasizing the need for further targeted research. This review provides a foundation for future investigations into JEV transmission in Australia, advocating for enhanced surveillance and standardized research methodologies to better understand and mitigate the virus’s impact.

## Introduction

Japanese encephalitis virus (JEV) is a mosquito-borne flavivirus responsible for the most common viral encephalitis in Asia, Japanese encephalitis (JE) ^1^. In humans, the virus primarily infects the central nervous system, which can result in symptoms such as fever, headache, vomiting, confusion, seizures and paralysis. In severe cases it can cause fatality, particularly in young children and older adults ^2^. An estimated 3 billion people live in areas at risk across Southeast Asia and the Western Pacific region, with an estimated mortality of 25 thousand deaths in 2015 ^1^. Beyond its impact on human health, JEV also carries significant economic implications, including reproductive losses in pigs and clinical encephalitis in horses during outbreaks ^3–5^. Combating JEV requires a One Health strategy, integrating human, animal, and environmental health efforts to address its complex transmission dynamics and reduce its societal and economic burden ^4^. The circulation of JEV is maintained in a complex transmission cycle between wildlife hosts and mosquito vectors, with subsequent spillover to domestic pigs and humans (Figure 1). Early investigations into the vertebrate hosts of JEV suggested two potential transmission cycles: a maintenance cycle between herons and mosquito vectors, and a spillover cycle between pigs, mosquitoes and humans ^6^. These early studies, undertaken in Japan between 1955 and 1960, included experimental infection studies on potential hosts, annual seroprevalence surveys, virus isolation attempts and vector feeding preference experiments ^6–8^. These studies identified herons as maintenance hosts of JEV because (i) their annual arrival coincided with new infections in swine and humans, (ii) they demonstrated high competence to infect vectors under experimental conditions, and (iii) under natural conditions they were preferentially fed upon by vectors. Pigs were identified as key spillover hosts due to strong evidence that they were infected before humans and after herons ^6^. While these studies have been foundational for our understanding of JEV transmission, the host roles identified in Japan are not representative in other areas where JEV is endemic or newly emerged. For example the role of pigs remains unclear; JEV has continued to infect humans in Singapore decades after the abolition of pig farming ^9^; in China, different genotypes of JEV were isolated in pigs compared to those in humans and mosquitoes ^10,11^; and in India and Bangladesh, despite low pig densities, recurrent epidemics of JEV still occur ^12^. Likewise, while the evidence supporting herons as hosts for JEV transmission in Japan is compelling, herons may be less important to transmission in other habitats, and many alternate avian species are yet to be investigated as JEV hosts. Transmission of JEV has been reported in more than 24 countries, many of which have not fully resolved the species involved in transmission and the role of different cycles to human and swine infections ^2^.

**Figure 1.**
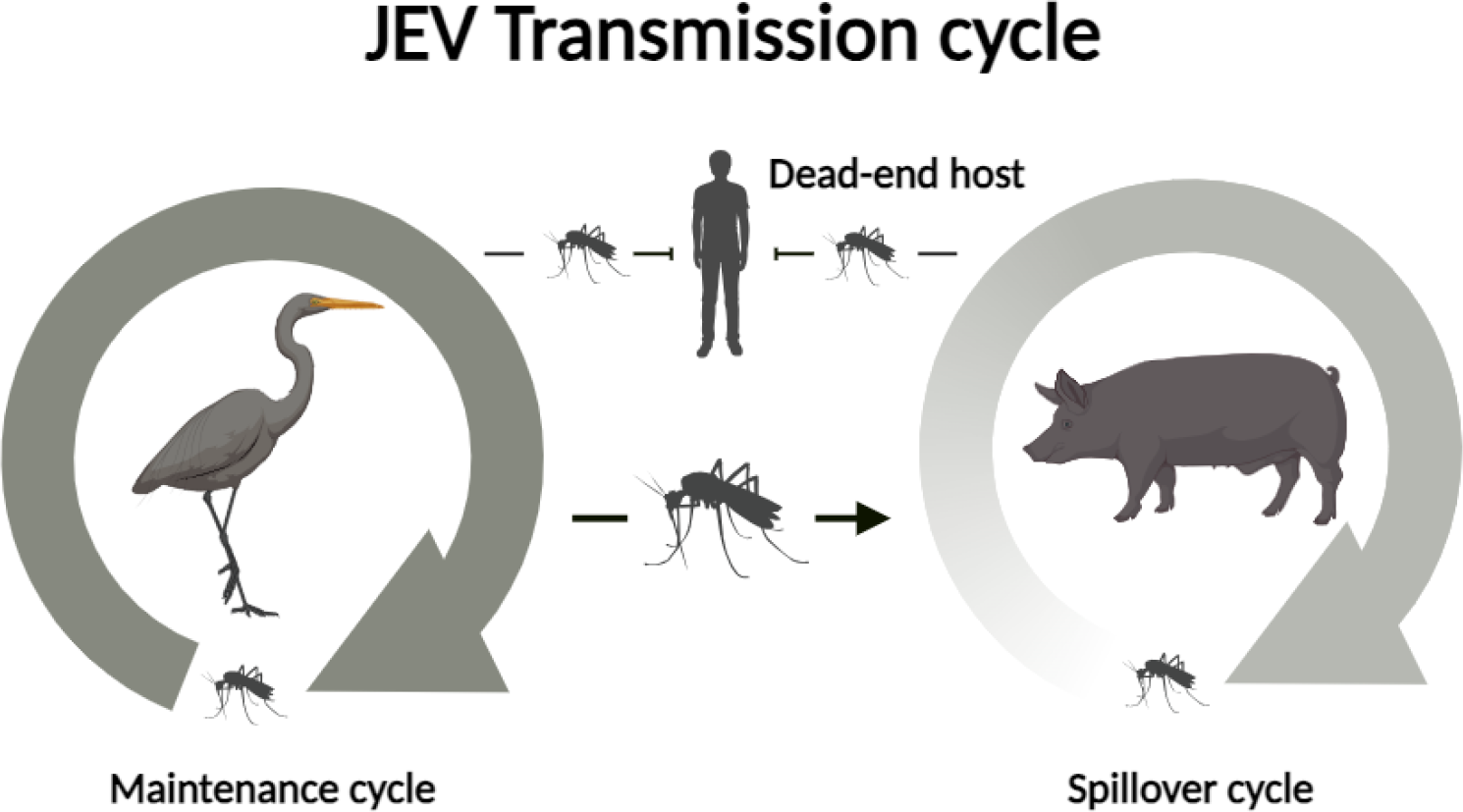
Transmission of JEV is dynamic with at least two different cycles. The maintenance cycle involves hosts that facilitate continued JEV transmission over time and between locations. Hosts in the spillover cycle become infected incidentally and sustain transmission for a limited time. Dead-end hosts (such as humans) do not contribute to JEV transmission.

Until recently JEV in Australia was primarily considered a risk for people travelling to endemic areas in Asia ^13^. Japanese encephalitis virus first emerged in Australia in 1995 when an outbreak occurred on an island in the Torres Strait (Figure 2) ^14^. Response to this outbreak was widespread and included the vaccination of 3,340 people, vector control and viral surveillance within domestic vertebrates ^14,15^. Natural transmission of JEV was not reported in Australia again for more than 25 years ^16^. In 2022, a large and widespread outbreak of JEV occurred across all but two states and territories in Australia (Figure 2) ^16^. This outbreak led to the deaths of seven people, infected over 80 piggeries and resulted in a loss of more than $3,000,000 AUD within the pig industry ^5,16^. The widespread nature of this outbreak raised questions about the introduction of JEV to Australia, the local hosts responsible for dispersal, and whether the virus would become established and lead to future outbreaks ^16^. These two different outbreak periods in Australia (1995-1998 and 2021-2022) are speculated to be associated with different transmission systems ^16^. The northern parts of Australia are hypothesised to maintain endemic transmission between highly abundant feral pigs and Culex mosquito species, though confirmation of this is highly challenging given the vast landscape, low human density and many challenges of trapping and testing feral animals. In southern Australia, transmission is hypothesised to be an epidemic system that arose due to opportunistic dispersion of migratory birds to inland regions affected by heavy rainfall and flooding associated with La Niña weather patterns ^17^. At present, there is not enough evidence to confirm or refute these theories.

**Figure 2.**
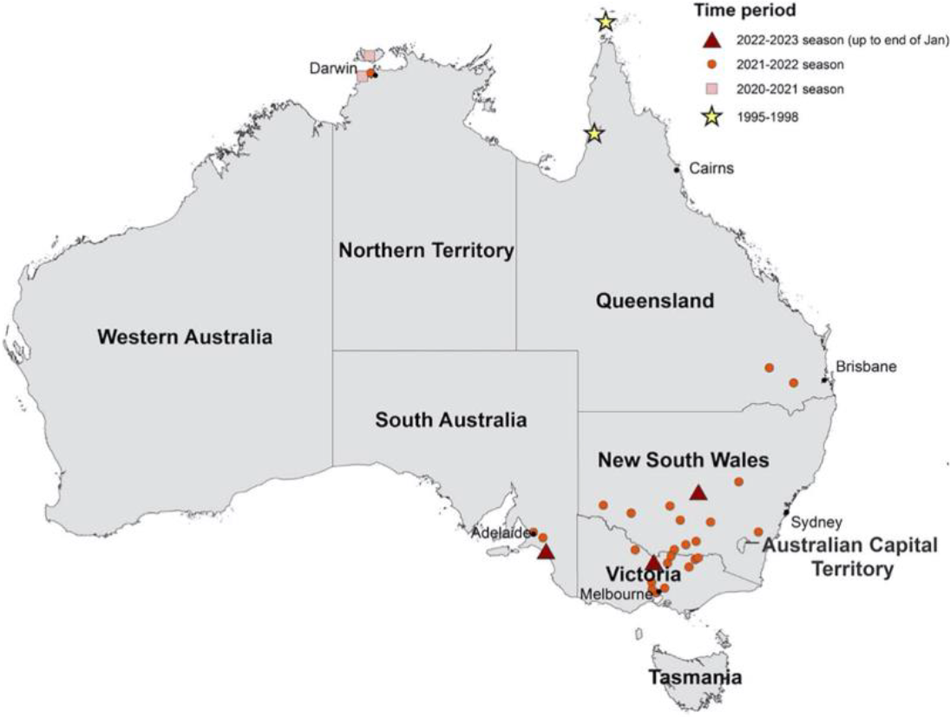
Residential location for 40 of 46 human JEV cases from the recent (2021-2023) outbreak and five of five cases of historical JEV outbreaks (1995 and 1998) in Australia. The location of symbols corresponds to the Local Government Area of residence of cases and may not reflect place of exposure. More than one case was reported for some LGAs. (Source: McGuinness et al 2023) ^18^.

The aims of this review were to identify the existing evidence for vertebrate hosts of JEV within Australia, collate data across studies, and identify gaps in our understanding of JEV transmission in Australia. Specifically, we synthesise data from studies on experimental infection, seroprevalence and virus isolation and calculate a host competence score for included species.

## Materials and Methods

We conducted a systematic review of the literature and extracted data from primary research studies. We included primary research studies of three data types: experimental infection, seroprevalence, and virus isolation studies. Experimental infection studies were included if they evaluated infection in native Australian species, migratory bird species that reside in Australia and imported or domestic species known to live in Australia (such as pigs, horses and water buffalo). Seroprevalence and virus isolation studies were included if they took place in Australia. A detailed search strategy and inclusion criteria can be found in Supplementary Methods, Textbox S1 and Table S1.

To compare results of experimental infection studies across species and observations, we calculated a metric of host competence building on an approach used in other studies ^19,20^. In summary, we calculated the host competence scores in the following way; first we calculated the mean duration of viraemia and mean peak viraemia across all individuals for each species from raw data extracted from the included experimental infection studies. Using these values, we generated a viraemia curve for each species by fitting a quadratic function and assuming that mean peak viraemia occurs halfway through the duration of viraemia (Supplementary Methods). From these viraemia curves, we calculated the area under the curve (AUC) to allow comparisons between species with high, but short lived, viraemias to those with low, long-lasting viraemias. Finally, we multiplied the AUC by the proportion of individuals of that species that developed a viraemia to account for variation in susceptibility within species. This calculation thus includes the mean peak and mean duration of viraemia within a species and considers the proportion of individuals that had a viraemic response. In doing so, it offers an estimate of a species’ capacity to infect susceptible vectors.

## Results

### Systematic review

We identified 20 studies that examined 22 vertebrate species present in Australia. Three studies were serosurveys and 17 were experimental infection studies. Most experimental infection studies took place outside of Australia (14/17; 82%). The three experimental infection studies conducted in Australia infected agile wallabies (*Macropus agilis*), black flying foxes (*Pteropus alecto*), brushtailed possums (*Trichosurus vulpecula*) Eastern grey kangaroos (*Macropus giganteus*), tammar wallabies (*Macropus eugenii*), Nankeen-night herons (*Nycticorax caledonicus*) and intermediate egrets (*Ardea intermedia*). Serosurveys tested for prior JEV infection in flying foxes, pigs and humans in Western Australia, the Northern Territory and the Torres Strait Islands. Humans and pigs were tested as part of outbreak investigations during the 1995 and 1998 JEV outbreaks in the Torres Strait Islands. We did not identify any studies that isolated virus from free-living vertebrates in mainland Australia, but one study isolated JEV from a sentinel pig in the Torres Strait in 1998 ^21^. A summary of all included studies is provided in Table S2.

### Seroprevalence studies

A total of 419 individuals from 11 species in Western Australia ^22^, the Northern Territory ^22^, the Torres Strait Islands and northern Queensland have been tested for antibodies (Ab) to JEV (Table 1) ^14,21^. Across the islands of the Torres Strait, seroprevalence was highest in pigs (*Sus scrofa*; 70%, 63/90), followed by horses (*Equus caballus*) (70%, 7/10), dogs (*Canis lupus familiaris*) (63%, 10/16) and chickens (*Gallus domesticus*) (0%, 0/6). In surveys conducted on the Cape York Peninsula in 1998, JEV Ab were found in 65% (13/20) of domestic pigs. However, none of the 113 feral pigs tested positive for JEV Ab, although 90 of the 113 exhibited cross-reacting flavivirus Ab ^21^. Surveillance of mega- and microchiroptera in Western Australia and the Northern Territory in 1998-1999 yielded potentially positive JEV Ab detections in black flying foxes ^22^. However, again, cross-reactivity within the flavivirus serological complex limits the meaningful interpretation of data from other bat species in this survey ^22^.

**Table 1.**
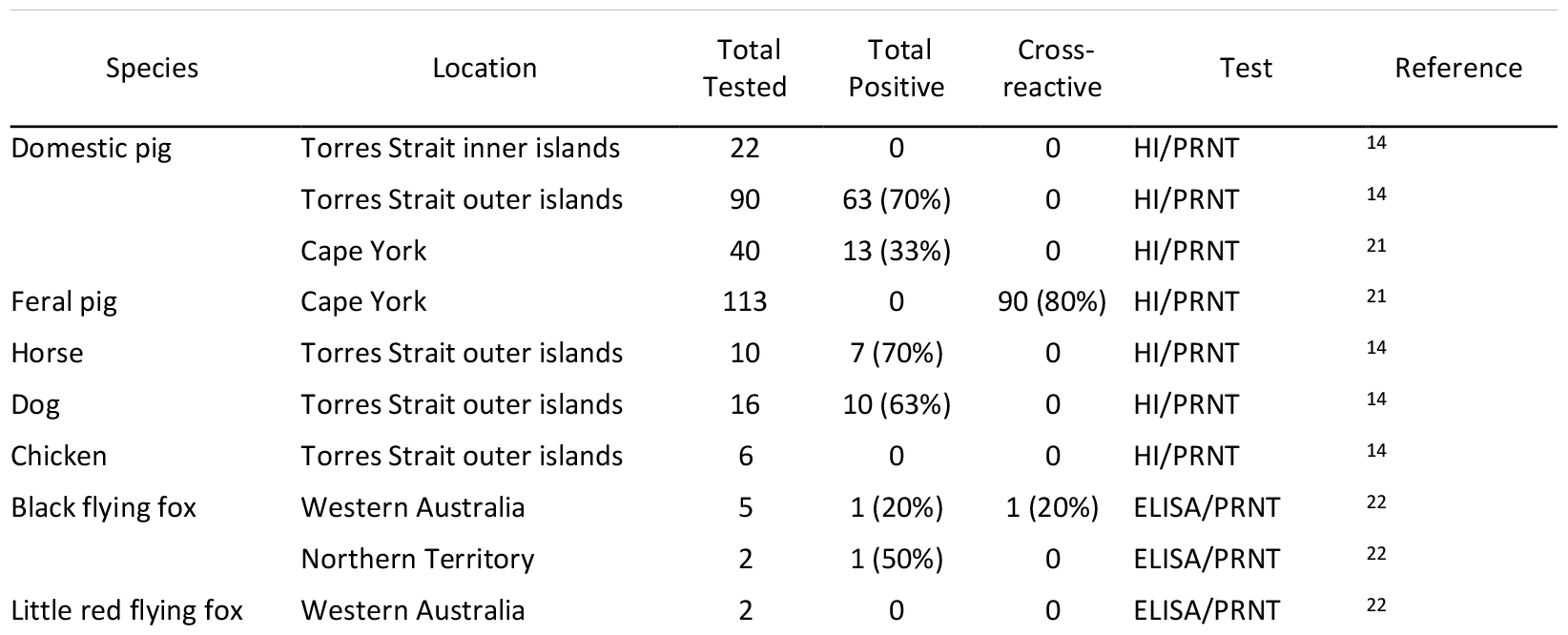
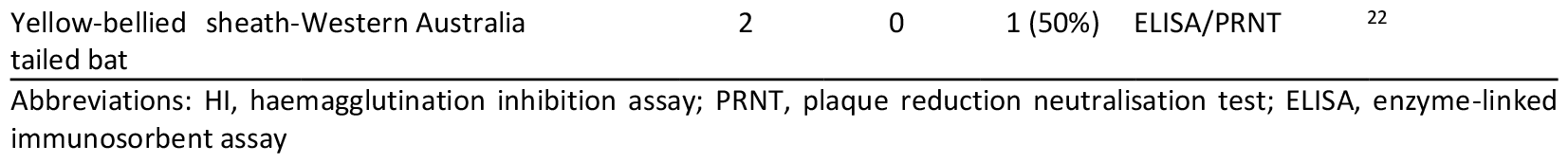
Summary of JEV seroprevalence studies conducted in Australia.

**Table 2.**
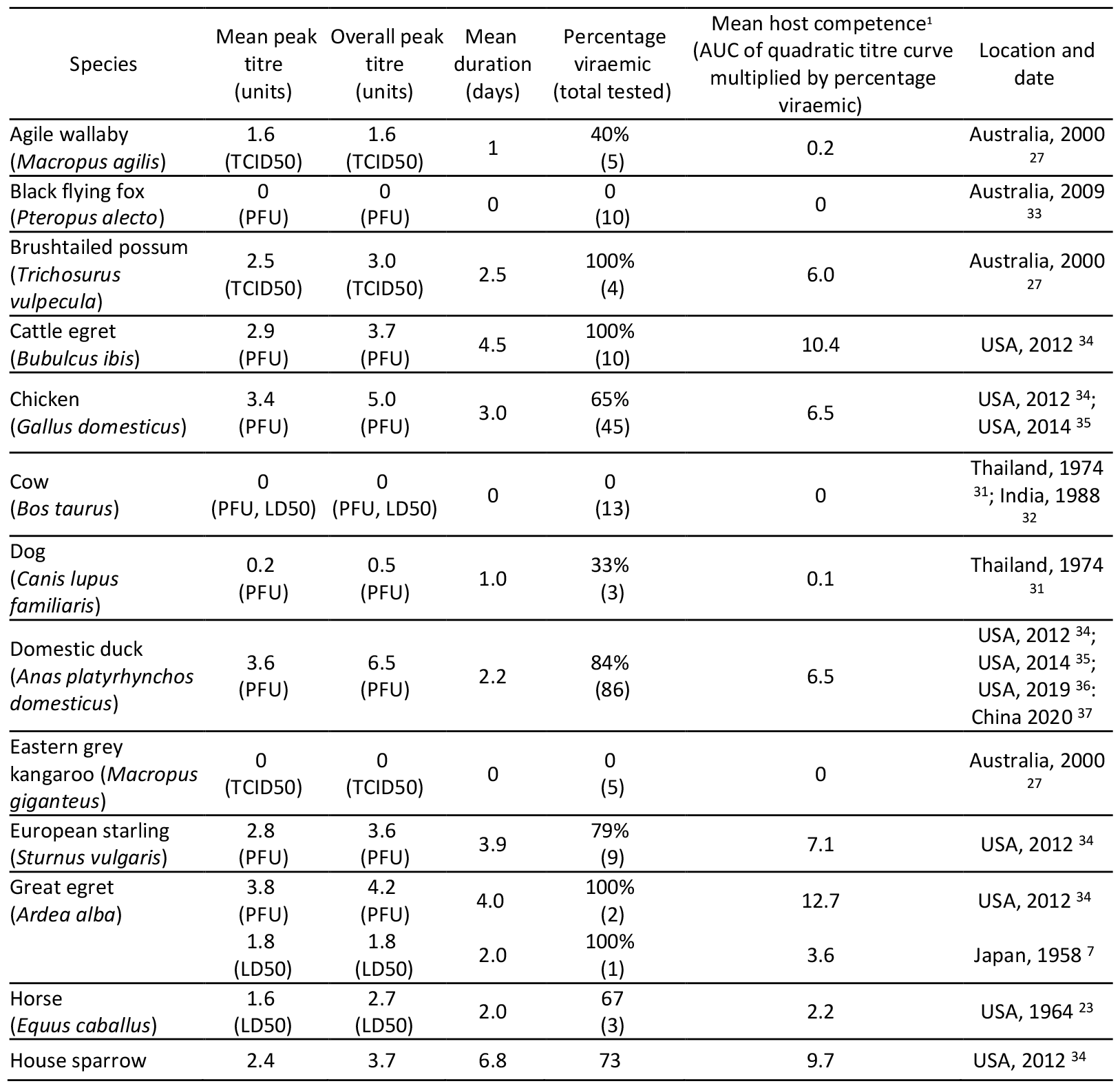
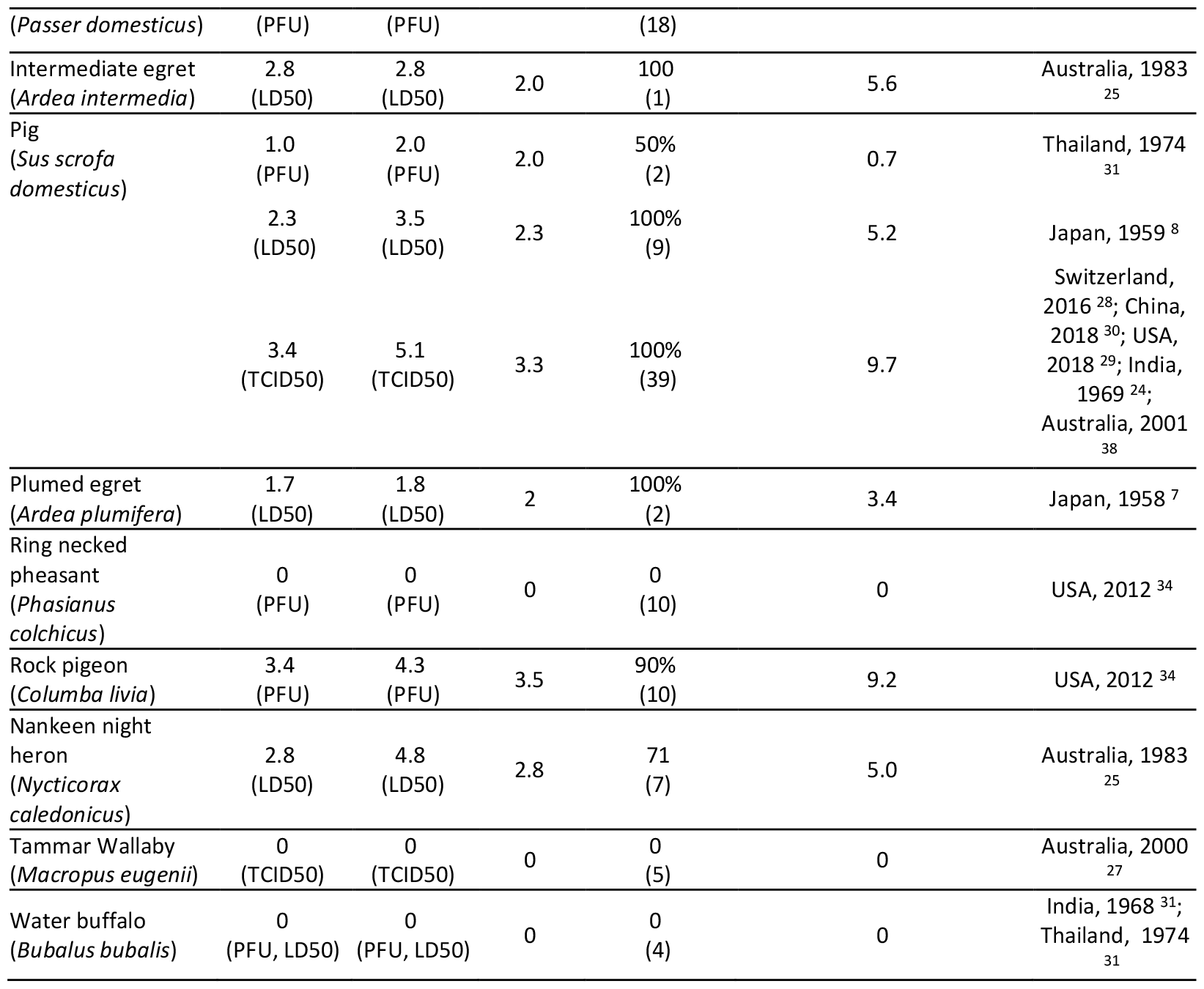
Summary of JEV experimental infection studies with Australian vertebrate species, and mean hosts competence calculation from experimental infection data.

### Viraemia from experimental infection

The 17 experimental infection studies included 22 Australian vertebrate species. All individuals were inoculated with strains of JEV from either the genotype 1 (GI), genotyoe 2 (GII) or genotype 3 (GIII) genotypes. The methods of inoculation and metrics used to measure viraemia differed between the studies. Studies prior to 1988 used LD50 ^7,23–26^ in mice to measure viraemia, while more recent studies used tissue culture infectious dose 50 (TCID50) ^27–30^ and plaque forming units (PFU) ^31–37^. Across all studies, the median sample size for each species was five, with ducks and chickens having the largest sample sizes of 48 and 41, respectively ^35^. A two-day old duckling had the highest reported PFU viraemia at 6.4 PFU with a duration of four days ^35^, while a seven-week old pig had the highest TCID50 measured viraemia at 5.1 TCID with a duration of three days ^28^. Nankeen night herons had the highest LD50 measured viraemia at 4.8 LD50 with a duration of 3 days. The species with the longest mean duration was the house sparrow (6.8 days). Cows (*Bos taurus*), water buffalo (*Bubalus bubalis*), tammar wallabies, Eastern grey kangaroos, ring necked pheasants (*Phasianus colchicus*) and black flying foxes did not produce a detectable viraemia; however black flying foxes were able to infect feeding mosquitoes with JEV ^33^.

### Host competence scores

Mean host competence (MHC) is the term we use to succinctly refer to the host competence calculation, i.e., the area under the curve (AUC) of the quadratic viraemia curve multiplied by the proportion viraemic. Mean host competence describes the average host competence across all individuals of a species. Please refer to the supplemental methods for a full description of this calculation.

All five Ardeidae species had high mean host competence scores, ranging from 3.4-5.6 for those measured with LD50 and from 10.4-12.7 for those measured with PFU (Figure 3). The great egret and cattle egret had the two highest mean host competence values across all species measured with PFU (10.4 and 12.7) and the intermediate egret had the highest mean host competence of those measured with LD50 (5.6) (Figure 3). The 27 pigs measured with TCID50 and LD50 had high mean host competence (6.9 and 5.2, respectively), while the two pigs measured with PFU returned a low mean host competence of 0.7 (Figure 3).

**Figure 3.**
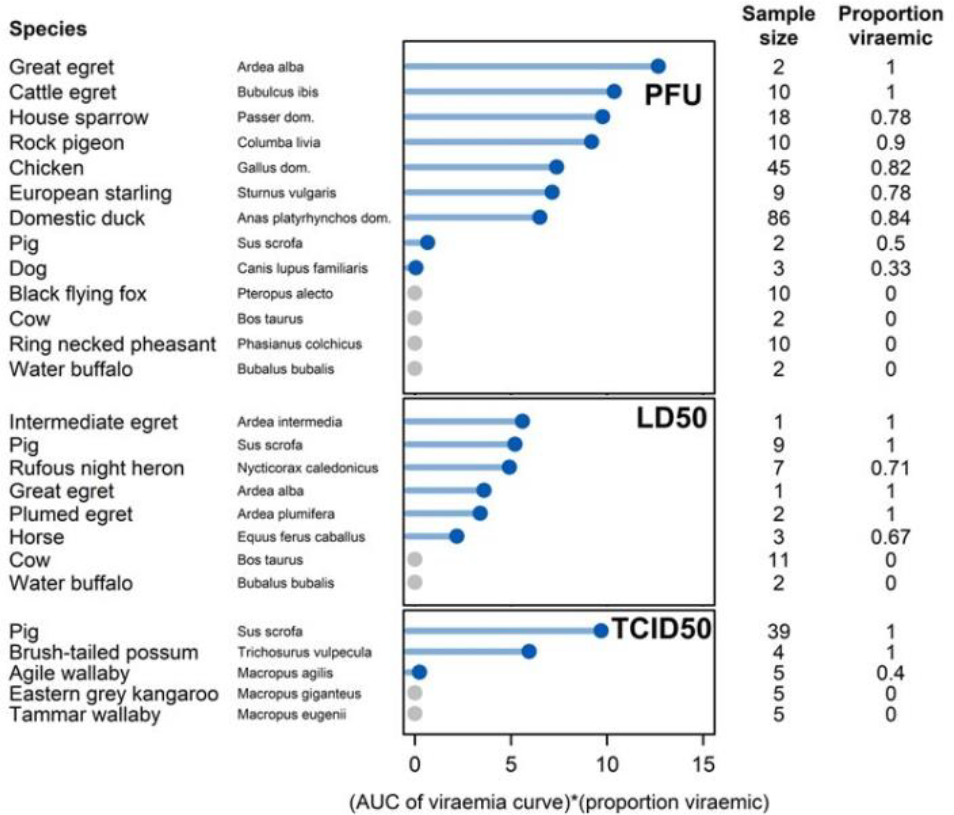
Mean host competence score, sample size and proportion viraemic for each species included in the review. Species are grouped by unit measure of viraemia (PFU LD50, TCID50) and ranked from highest to lowest mean host competence score (solid circles). Abbreviations: PFU, plaque-forming units; TCID50, tissue culture infectious dose 50; LD50, lethal dose 50

Three other bird species, the rock pigeon (*Columba livia*; 9.2), European starling (*Sturnus vulagaris*; 7.1), and house sparrow (*Passer domesticus*; 9.7), have mean host competence scores similar to the two Ardeidae species included in the same study (great egret [*Ardea alba*; 12.7] and cattle egret [*Bubulcus ibis*; 10.4]) (Figure 3) ^34^[32]. The agile wallaby (*Macropus agilis*; 0.2), dog (*Canus lupus familiaris*; 0.1), and horse have low mean host competence scores attributed to low titres and short durations of viraemia (Figure 3).

## Discussion

Japanese encephalitis virus is sustained in the environment by the interactions between a multiple vector and vertebrate host species ^39^. Not all vertebrate host species contribute equally to transmission with some species considered more broadly as maintenance hosts, spillover hosts or dead-end hosts ^40^. Vertebrate host species integral to the persistence of JEV – whereby their absence would lead to a reduction or elimination of JEV within the community – are considered maintenance hosts. In contrast, vertebrate species that contribute to JEV transmission, but only for short periods of time or with repeated introduction are considered spillover hosts ^41^. Those species which cannot infect susceptible vectors are considered dead-end hosts. In the Australian context the role of different species as maintenance, spillover or dead-end hosts is unknown but has important implications for the management and mitigation of future transmission. Here we synthesise our findings of Australian species in the context of these different species roles but acknowledge that understandings of JEV transmission in Australia is incomplete and with new data these potential species roles could change.

### Possible maintenance hosts of JEV in Australia

In endemic areas of JEV transmission (such as Japan), the Ardeidae family of birds have long been considered the key maintenance hosts of JEV because they generate high, long-lasting viraemias, are frequently and preferentially fed upon by vectors, have regular population turnover, and consistently demonstrate JEV exposure in free-living populations ^6–8^. In Australia, there are twelve species of the Ardeidae family, of which experimental infection data was available for five species ^7,25,34^. These five species had consistently high mean host competence scores (3.4-12.7) compared to other non-Ardeidae species experimentally infected with JEV ^7,25,34^. Additionally, there is strong ecological support for Ardeidae species as maintenance hosts based on their high abundance and colonial breeding in areas of JEV transmission in Australia, and their wide distribution across the country ^17^.

While the host competence and ecology of Ardeidae species provides strong evidence for them as potential maintenance hosts of JEV in the Australian landscape, there is limited evidence demonstrating the interactions between vectors, Ardeidae species and JEV. In a meta-analysis of all bloodmeal studies in Australia, only one *Culex annulirostris* (an important JEV vector in Australia) was reported to have fed on Ardeidae ^42^. Yet, this species is also considered the primary hosts of Murray Valley encephalitis virus (MVEV), a flavivirus closely related to JEV, which suggests vectors are frequently interacting with Ardeidae species. However, this adds more complexity to the potential role of Ardeidae species as JEV hosts because cross-immunity from MVEV could exclude certain populations from the JEV transmission cycle. More studies are needed to identify previous JEV exposure in Ardeidae populations, mosquito feeding patterns across space and time, and the immunological implications of previous MVEV infection. Without these studies, our understanding of Ardeidae species in past or future JEV outbreaks is incomplete.

Non-Ardeidae bird species have been considered maintenance hosts of JEV in a number of other studies ^6,36,43^. The original investigations of JEV hosts in Japan noted that although Ardeidae species were their primary focus, this was not intended to exclude other species from consideration in JEV transmission ^8^. In fact, experimental infection studies in the last decade identified high viraemias in rock pigeons, house sparrows, and European starlings ^34^. When comparing host competence scores calculated in this review, these three species have scores below the Ardeids and higher than pigs in their capacity to infect susceptible vectors. There is evidence of natural exposure to JEV in these three species in countries where JEV transmission is endemic. Specifically, in wild pigeons in India, JEV Ab were prevalent throughout the year ^44^. In Thailand, a small percentage of wild tree sparrows had Ab to JEV ^31^. And in Japan, 20-37% of wild-caught sparrows had Ab to JEV ^45^. The ecology of these species also supports their potential as maintenance hosts; they congregate in high numbers, are short-lived and have moderate clutch sizes allowing for the availability of a susceptible population ^46–48^. In Australia, these species have not yet been investigated as potential maintenance hosts, and there is an absence of data on mosquito feeding patterns for these species, as well as previous exposure to JEV.

Flying foxes (genus *Pteropus*) have the potential to be maintenance hosts because (i) they are capable of infecting mosquitoes with JEV ^33^, (ii) there is some evidence to suggest prior exposure to JEV in Australia ^22^ and (iii) Australian JEV vectors are known to feed on them (which confirms a potential transmission pathway) ^19^. Recent investigations of JEV transmission in Indonesia (where the same genotype of JEV that caused the outbreak in Australia circulates endemically - genotype 4 [GIV]) found 5.6% (n = 373; from 22 species) of bats sampled were positive for JEV using RT-PCR ^49^, while another study found that 4.2% (3/70) of Pteropus sp. In Indonesia had been previously exposed to JEV ^50^. These studies indicate that JEV can be transmitted from mosquitoes to bats but does not confirm that bats are transmitting the virus further. The evidence in Australia is mixed, the only seroprevalence study estimated that <3% (3/119) of bats had prior exposure to JEV, but the authors note a high potential for cross-reactivity to other closely related flaviviruses ^22^. An Australian experimental infection study found no detectable viraemia in any of the ten black flying foxes included, but at least three of these individuals were able to infect susceptible Cx. annulirostris vectors ^33^. Bats are known to have unique immune systems, which has been observed in the experimental infection of JEV in Microchiropteran species ^51^. Tricoloured bats (Perimyotis subflavus; native to North America) were capable of maintaining viraemia after induced hibernation periods and were also observed to be viraemic after feeding on infected vectors ^51^. Australia is home to over 90 species of bats that inhabit diverse ecological niches, travel large distances, and gather in dense communities ^52^. While there is a potential that some of these species could maintain JEV transmission, more evidence is needed to identify which species and their full capacity to do so.

Australia has a feral pig population of over 24 million distributed across at least 45 percent of the continent ^53^. This large and widely distributed population is conducive to participation in the maintenance cycle, as opposed to domestic pigs, which live in isolated populations with minimal to no movement. Also, some of these feral pig populations co-occur with JEV vectors that are active year-round in northern Australia ^19^. Feral pigs accounted for 82% of vector bloodmeals identified (in the absence of domestic pigs) at the location that yielded the first Australian mainland isolate of JEV from mosquitoes ^54^. Wild boar sera were collected in Japan and 86% were positive for neutralizing Ab against JEV, even in winter ^55^. The prevalence of JEV Ab in wild boar has been cited as 83% in Japan ^56^, 66% in South Korea ^57^, 32% in Indonesia ^58^ and 15% in Singapore ^9^. In March of 2022, as part of a routine animal health survey, a small number of feral pigs in northern Australia tested positive for JE ^59^. And retrospective serology from 2020-2021 shows evidence of exposure in three feral pigs in northern Australia ^59^.

Data from blood meal analyses have also found support for vector-pig interactions. All 14 confirmed JEV vector species, eight of which are present in Australia, are known to feed on feral pigs ^60^. The hypothesised primary vector of JEV in Australia, *Cx. annulirostris*, has been found to predominantly feed on feral pigs in a location with JEV activity ^54^. This finding is not consistent across all locations, but it points to a possible transmission route. In general, feral pigs are not the primary bloodmeal source for JEV vectors in Australia, even in areas with feral and domestic pig populations ^19^. However, JEV has been reported to spread even in areas with low porcine feeding rates, as demonstrated by Hall-Mendelin et al in northern Australia which could suggest the importance of other hosts in the transmission cycle ^54^. Theoretically, feral pigs in northern Australia could participate in a maintenance transmission cycle, but this prospect is yet to be adequately investigated.

### Possible spillover hosts for JEV in Australia

A JEV spillover host is a species that can transmit the virus, but whose biological/ecological characteristics limit their ability to independently maintain transmission (i.e. in the absence of other host species). In this context, we consider domestic pigs and brushtail possums to be potential spillover hosts in Australia. Australian domestic pigs are possible spillover hosts because, along with their strong host competence (mea of 5.2 [LD50], 6.9 [TCID50] and 0.7 [PFU]), they were only seasonally infected in areas with endemic JEV transmission, with summer months coinciding with higher infection rates. Additionally, transmission to domestic pigs does not persist year to year in Australia ^61^. During the 2021/2022 outbreak in Australia, similar observations were made where over 80 domestic pig farms had detected JEV within the span of three months ^5^ and yet despite increased surveillance was not detected at any Australian piggery the following year ^5^. The inability for domestic pigs to sustain transmission between years, irrespective of the whether this is due to their immunology, local ecology or piggery practices, supports the hypothesis that they are a spillover host. In Asia, where JEV is endemic, domestic pigs can also be amplifying hosts ^43,61^. An amplifying host increases the level or circulating virus, which leads to pathogen pressure on humans ^39^. In many locations across Asia, domestic pigs fit this definition because they live in dense populations near humans and develop high viraemias ^61^. After the 1995 outbreak in the Torres Strait, a clear link was made between the proximity of domestic pigs and *Cx. annulirostris* breeding sites that could have amplified infections in humans ^14^. During the 1998 outbreak in the Torres Strait and Northern Peninsula Area (NPA) of Cape York, however, the relationship between domestic pigs and human cases was less clear ^21^. A successful vaccination campaign in the Torres Strait after the 1995 outbreak, along with unique climate factors and a lower percentage of NPA households keeping pigs (10% in NPA compared to 50% on Badu Island in the Torres Strait) likely contributed to reduced transmission to humans in these areas ^15,21^. In mainland Australia, commercial pig farms are usually located away from human population centres. During the 2021/22 outbreak in Australia, the 45 human infections were hypothesized to be from the maintenance cycle of JEV (rather than from proximity to domestic pigs) because a number of cases were not in close proximity to infected piggeries.

Overall, domestic pigs in Australia are competent hosts, they are infected when JEV is circulating, and have the potential to increase pathogen pressure on humans in situations where they are kept in proximity to communities. The link between the maintenance cycle of JEV and domestic pigs has not been identified, and the degree to which domestic pigs increase the risk of JEV transmission to Australians requires further evaluation.

### Species with unknown roles

Not enough data is available to distinguish the potential role of domestic ducks and chickens as maintenance or spillover hosts. Although the host competence scores for ducks indicate their potential as hosts (6.5 MHC), there are two caveats that must be considered. First, the individuals identified in this review are domestic ducks (*Anas platyrhynchos domesticus*), and their results may not extrapolate to other wild species in the genus Anas. Second, the majority of the individuals in the original experimental infections were newly hatched (less than three weeks old; 60/86, 70%) and, as Cleton et al. (2014) demonstrated, viraemia is a function of age in young ducks, where younger ducks (<20 days) generate a much higher viraemic response than older ducks ^35^. Within our dataset, the MHC for ducks over the age of 20 days is 1.9; the MHC for ducks under the age of 20 days is 9.2. This difference indicates that adult domestic ducks might be less competent hosts than young ducks (and less competent than other bird species), but it is not clear how this would affect the ecology of JEV transmission.

The role of domestic ducks as either maintenance or spillover hosts therefore remains unknown. On one hand, in JEV endemic regions, both wild and domestic duck species are frequently infected. In seroprevalence studies, significant numbers of ducks were found to have Ab to JEV in India (*Anas clypeata, Anas crecca, Aythya fuligula, Anas strepera*) ^44,62^ and Thailand (*Anas platyrhynchos domesticus*) ^31^. In Indonesia, Ab against JEV were found in ducks (20.6%), with no difference in seroprevalence between domestic ducks kept closely with pigs compared to those reared without pigs ^58^. In Japan, 85.9% of wild ducks were positive for JEV Ab (*Anas poecilorhyncha, Anas platyrhynchos, Ana acuta, Anas Penelope*) ^63^. On the other hand, unlike pigs, the epidemiological significance of domestic ducks living in proximity to humans has not been studied ^64^. Death associated with natural JEV infection is not observed in duck farms, and therefore JEV outbreaks might be ignored ^37^.

Like domestic ducks, chickens have been successfully used in JEV mosquito transmission studies, suggesting chickens may exhibit sufficient viraemia and receptivity to infection to participate in natural transmission cycle ^43^. However, in an experimental infection study in 1951, chickens were deemed relatively unsusceptible, with the virus detected in only 4/12 (33%) at low titres ^65^ (data not included in host competency calculations because of incompatible methods). Overall, chickens have a strong MHC of 6.5 but only 65% (34/52) of individuals developed a viraemia, and all of these viraemic chickens were young individuals, illustrating that viraemia is also a function of young age in chickens ^35^.

### Dead-end hosts

Eight of the species included in this review developed no more than a trace viraemia and, unlike flying foxes, did not demonstrate the ability to infect mosquitoes. In the experimental infection study with macropods, two agile wallabies and two tammar wallabies developed trace viraemias, and one Eastern grey kangaroo generated a low-level Ab response ^27^. The lack of viraemia generated by these three species indicates that they are not likely to contribute to transmission, however more replication is needed to confirm. Furthermore, in mosquito feeding studies across Australia, macropods make up a large proportion of bloodmeals in JEV vectors, including Cx. annulirostris and Cx. sitiens ^19^. This suggests that there is a possibility that if infected vectors were to feed on susceptible macropods, the macropods could play a role as dilution hosts ^66^. However, exhaustive evaluation of this hypothesis would require studies on transmission pathways and exposure before it could be confirmed. It is plausible that other macropods species differ in their response to JEV, given that macropods play a role in other arboviral transmission in Australia ^67,68^.

In endemic areas of JEV transmission, horses have often been considered dead-end hosts alongside humans ^69^. However, closer inspection of this information reveals limited and conflicting data. A single experimental infection study, with a small sample size (n=3), found that horses developed a low mean peak viraemia (1.6 LD50) ^23^. In the same study, Gould et al. (1964) demonstrated experimental horse-to-chick and horse-to-horse transmission via mosquitoes ^23^. Horses do suffer morbidity from JEV infection with fatality reported between 5-30% ^69^, and infections in unvaccinated horses as high as 73% ^70^. Consequently, countries such as China, Japan, Korea, and India practice JEV vaccination of their horses, with the result that equine infections are now rare and predominantly sub-clinical in these countries ^3,71^. However, horses, unlike pigs, do not appear to significantly contribute to JEV transmission to mosquitoes, and their smaller population size, slow turnover, and extended lifespan likely limit their role as hosts ^72^.

The prevailing theory that cattle are dead-end hosts is supported by experimental infection studies showing no detectable viraemia in inoculated cattle ^26,31^. Several studies report high seroprevalence (21% - 51%) among cattle in Asia ^73–75^, indicating that JEV is transmitted to cattle and that they generate an immune response. Boyer et al (2021) found that in Cambodia, JEV vectors demonstrated a preference to cattle over chickens and humans, with pigs as the secondary choice ^76^. Overall, cattle do not demonstrate disease or generate a viraemia, and therefore are most likely dead-end hosts. Similarly, to macropods, a high proportion of bloodmeals in northern Australia came from cattle suggesting that they too could play a role as dilution hosts depending on the transmission and population dynamics of both vectors and cattle ^42^.

Several other species of free-ranging vertebrates (water buffalo, dogs, and ring-necked pheasants) were included in this review and are considered dead-end hosts for JEV transmission. Although these species may become infected with JEV and demonstrate a high prevalence of Ab, they develop low viraemic responses ^77^. A small number of water buffalo (n=2) were experimentally infected and neither developed a viraemia ^24^. A single experimental infection study on dogs failed to detect more than a trace viraemia in one of the three (33%) of the infected individuals. In Cambodia, JEV seroprevalence was 35% in dogs ^78^ and a Malaysian study showed that 80% of dogs had JEV Ab (commercial IgG ELISA) ^74^. While rats and mice experimentally develop HI Ab to JEV ^79^ and 45.8% of rats sampled in south China were positive for JEV-reactive IgG Ab ^80^, rodents are not known to play a role in transmission of JEV in Asia ^80^. Snakes have rarely demonstrated viraemia, however some species show high prevalence of JEV Ab ^81–83^. Interestingly, ring-necked pheasants were the only bird species in this review that did not generate a viraemia ^34^. Nemeth et al. remark that two species of the order Galliformes (i.e. chickens and ring-necked pheasants) generate low to undetectable viraemia ^34^. Of note, humans are not considered spillover hosts of JEV because, unlike with other arboviruses, humans do not contribute to onward transmission; each human infection is considered an epizootic spillover event ^39^.

### Limitations and future research

This review is limited by the scarcity of research on Australian hosts and the important work that needs to be done. Specifically, the limited number of species studied to date presents a major knowledge gap for the hosts of JEV in Australia, especially when considering the diversity of Australian species across habitats and climates. Additionally, we could not directly compare the physiological competence of species due to inconsistent methods, poor reporting, and small sample sizes. Each studied varied in the way they measured virus titres, even when using the same general approach.

Experimental infection studies aim to illuminate one step in the transmission cycle (host viraemia) and interpretation of their results should not include assumptions about subsequent steps. Most experimental infection studies in this review, for example, did not expose viraemic hosts to feeding mosquitoes to determine onward transmission. Furthermore, some mosquito vectors are specialist feeders, and although a vertebrate can develop a high viraemia they may not be fed upon by the competent JEV vector species. With regards to the Australian seroprevalence surveys, certain species were more likely to be targeted than others. The surveys that took place after the 1995 and 1998 outbreaks in northern Australia focused on the communities that were affected and did not contend with the idea of more widespread transmission of JEV. They did not sample the wild bird population, marsupials, or bats. The lone Australian seroprevalence survey of bats was not randomly sampled bats and limited by cross-reactivity with other flaviviruses.

The experimental infection studies in this review used strains of JEV from the GI, GII or GIII genotypes. Most studies used only one strain of JEV, some used multiple strains of the same genotype, and only one study used two genotypes in the same study. Xiao et al used the same methods to experimentally infect piglets with either GI or GIII ^30^. They found little difference between the genotypes, with a piglet inoculated with G1 producing the highest viraemia. The pooled results from all experimentally infected piglets would indicate that GIII produces a higher viraemic response in piglets. The JEV GIV genotype caused the Australian outbreak in 2021/22 – no experimental infection studies have been conducted with GIV. Therefore, we caution labelling one genotype more transmissible than another because the differences between species and genotypes could be due to experimental design or other biases.

Australian mosquito species likewise differ in their ability to transmit JEV, with each species varying in its host-feeding patterns, vector competence, and population dynamics, among other factors ^19^. A review by van den Hurk et al demonstrated that *Cx. annulirostris* is the primary vector species in Australia ^19^. Along with pigs and birds, *Cx. annulirostris* readily feed on marsupials, humans, and other placental mammals, a generalist feeding strategy that could result in a variety of host species being exposed to JEV ^19^. Future studies should investigate vector-host interactions across space and time, particularly at the interface between piggeries and wild bird populations ^19^.

## 6. Conclusions

This review set out to synthesise the existing evidence on Australian vertebrate hosts of JEV. Our findings suggest that domestic pigs (particularly during the 2021/2022 outbreak) and feral pigs, may play roles in the spillover cycle. Ardeidae birds and other non-Ardeidae bird species, along with flying foxes, emerged as potential maintenance hosts, but the absence of comprehensive data on their interaction with vectors and prior JEV exposure hinders a conclusive determination. The roles of domestic ducks, chickens, and other groups such as macropods, horses, and cattle are less clear, with some evidence pointing towards their designation as dead-end hosts. The complexity of host-virus-vector interactions, influenced by ecological, biological, and environmental factors, underscore the challenge of identifying definitive host roles.

In conclusion, our systematic review highlights significant gaps in the current understanding of the host roles in the Australian JEV transmission cycle. Despite the identification of 22 Australian vertebrate species in experimental infection studies, the lack of comprehensive ecological context limits definitive conclusions about their roles as maintenance or spillover hosts. Future research should focus on expanding the range of species studied, particularly marsupials and birds, with standardized methodologies to facilitate accurate comparisons. Enhanced JEV surveillance, including year-round monitoring, especially in northern Australia, is essential to understand the virus’s maintenance and introduction patterns. Only through such comprehensive and integrated research efforts can we hope to fully elucidate the transmission dynamics of JEV and develop effective strategies for its control and management in Australia.

## Author Contributions

Conceptualization, E.S., B.J.T., M.M., H.I.M and K.T.M.; methodology, M.M, E.S. and K.T.M; formal analysis, M.M., B.L., T.A., E.S., and K.T.M.; writing—original draft preparation, K.T.M.; writing—review and editing, B.L., M.M., T.A., B.J.T., E.S, H.I.M, and K.T.M. All authors have read and agreed to the published version of the manuscript.

## Acknowledgements

The authors thank David Williams for sharing unpublished data, and Stacey Lynch for useful discussions about this review.

## Data Availability

Statement: The code and data is available via Github at https://github.com/ThaboMoore/JEV-Host-Review.git.

