## Supplementary Methods for "Australian vertebrate hosts of Japanese encephalitis virus; a review of the evidence"

**Supplementary Materials**

**Supplementary Methods**

*Data collection and extraction*

We searched for primary research papers that used experimental infection, seroprevalence, or virus isolation methods from 1950-2022. First, we searched Web of Science (WoS; March 2022) for articles using the search terms in Textbox S1. Then, we searched bibliographies for additional references. Review studies, studies involving co-infections with multiple viruses, and studies that did not include Australian species or species that reside in Australia were excluded. We considered species relevant to an Australian JEV context if they reproduced in or migrated through Australia. We included experimental infection studies conducted in other countries, but we required seroprevalence and virus isolation studies to be conducted in Australia. We reported our results in line with the Preferred Reporting Items for Systematic Reviews and Meta-analyses (PRISMA), which has been described for use by researchers conducting systematic reviews. A PRISMA flow chart showing the article selection process is presented in Figure S1. A list of included publications is provided in Table S2.

KTM exported the search results, removed duplicates, screened all titles and abstracts for inclusion, and reviewed eligible full-text articles against the inclusion and exclusion criteria (Table S1). TSA, BL, and MJM each screened 30-50 abstracts (and full texts where the abstract indicated potential for inclusion). A senior researcher (ES) screened a random sample of 30 abstracts (and full texts where the abstract indicated potential for inclusion) and confirmed the consistent application of the inclusion and exclusion criteria. ES also provided guidance when eligibility based on a full-text review was unclear. Reasons for exclusion were recorded for all excluded studies (Figure S1).

For each article we extracted the following information: author, year of publication, species (common and scientific name), number of individuals, the dose of JEV inoculation (and unit of measurement), method of inoculation, viraemia measurements, the proportion of individuals that tested positive for antibodies to JEV, and the strain of JEV used (and source of strain). Where individual infection data was only available from the published figures and not from tables, we extracted raw titre values at each day post-inoculation from figures using WebPlotDigitizer v4.5 (Rohatgi 2021) and subsequently summarised into target metrics for each experimental group.

Raw viraemia data was provided by studies in one of two forms: summarised or daily. Studies that presented summarised viraemia data provided the duration of viraemia for each individual, the peak viraemia over that duration and the number of individuals that did not generate a viraemia. Studies that presented daily viraemia data listed the daily titre value for each individual, from which we determined the duration, peak viraemia and proportion viraemic. Studies were excluded if they provided a range of durations for a group of individuals or if the viraemia data was averaged from a group of individuals.

*Host competence calculations*

Data was compiled and analysed using RStudio (version 2023.06.0) and packages 'dplyr', 'tidyr', 'scales', and 'pracma'.

Initially, data were loaded and cleaned using the dplyr and tidyr packages. The dataset was bifurcated into two categories: 'days-post-infection' (DPI) data and summarized data. DPI data were summarized for each experimental replicate (individual animal), focusing on peak titre and duration of viraemia. These data were then grouped and consolidated by unique species and titre unit combinations, enabling calculation of average and maximum peak titres, average viraemia duration, and the number of viraemic individuals per species.

For the viraemia data, quadratic approximation was applied to model the viraemia curves. The x-axis values were as follows: lower limit set to zero; the centre of the curve was set to the average duration plus one, all divided by two; the upper limit was set to average duration plus one. The y values were as follows: lower limit set to zero, the mean peak titre, and upper limit set to zero. Using the pracma package, the area under the curve (AUC) for each species was then computed. The AUC values were then multiplied by the proportion of viraemic individuals for each species.

**Supplementary Textbox 1.** Search terms used in Web of Science search.

AB= (((japanese AND encephalitis) OR JE OR JEV) AND (host OR reservoir OR wild* OR feral OR captive) AND (((experimental AND infection) OR (viraemia or viremia) OR (competence)) OR (virus AND isolat*) OR sero*))

**Supplementary Table 1.** Inclusion and exclusion criteria guiding study selection.

| **Inclusion criteria** |
|---|
| • Primary research<br>• Experimental infection with JEV OR seroprevalence of JEV antibodies OR isolation of JEV from individuals<br>• Non-human species<br>• Species that live, or have ever lived, in Australia<br>• Seroprevalence studies must be conducted on individuals living in Australia<br>• Experimental infection studies must report viral titre |
| **Exclusion criteria** |
| • Review study<br>• Experimental co-infection with multiple viruses<br>• Seroprevalence of flavivirus antibodies without specific to JEV (reword)<br>• Non-Australian species<br>• Viraemia measured via PCR |

**Supplementary Figure 1.** Preferred Reporting Items for Systematic Reviews and Meta-Analyses flowchart of study selection process.

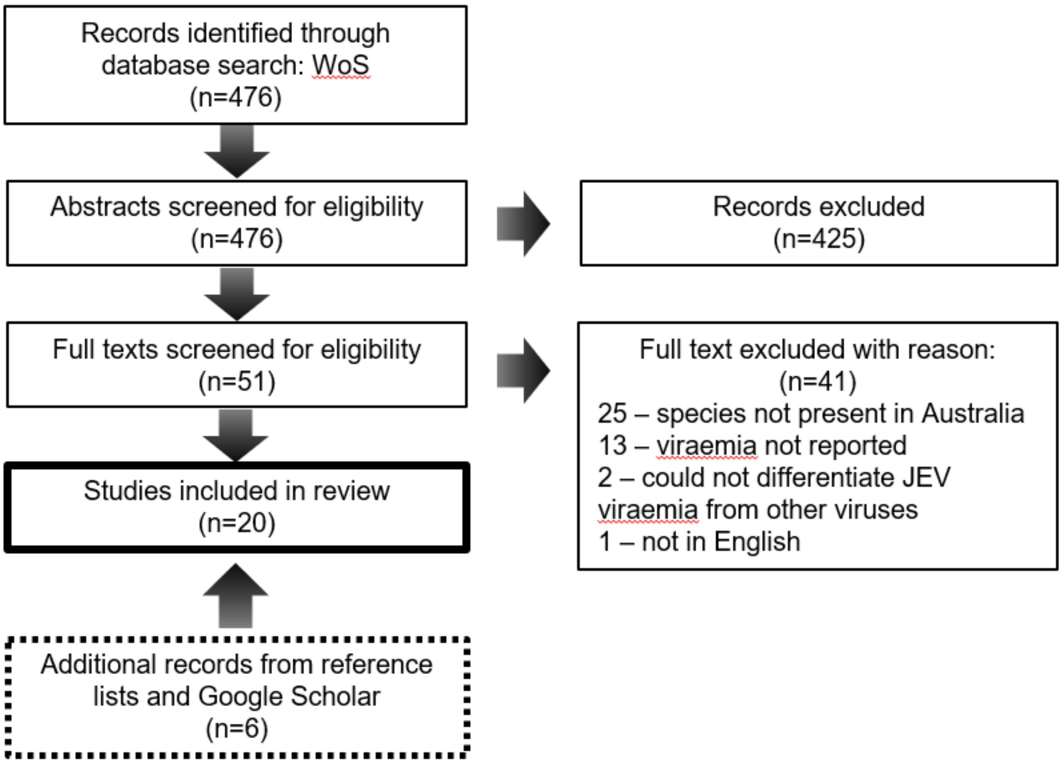


**Supplementary Table 2.** Description of included studies.

| Author | Strain | Method of inoculation | Country | Year | Measurement type | Sample size | Attempted transmission to mosquitoes? |
|---|---|---|---|---|---|---|---|
| Boyle | Nakayama (human origin) | Subcutaneous | Australia | 1983 | LD50 | 8 | No |
| Notes: authors do not list the volume (ml) used for LD50 calculation. Authors did not provide individual viraemia data, but the way the data was presented in Table 2 was sufficient to infer individual data. | | | | | | | |
| Carey | 9215 (mosquito origin) | Subcutaneous | India | 1969 | LD50 | 4 | Yes |
| Notes: we excluded on pig that was exposed to JEV by feeding mosquitoes, as we could not confirm that the mosquitoes were infected with JEV. | | | | | | | |
| Daniels | Isolate 4054 | Subcutaneous | Australia | 2000 | TCID50 | 20 | No |
| Gould | FM380 (mosquito origin) | Mosquito | US | 1964 | LD50 | 3 | Yes |
| Gresser | M5/596 (mosquito origin) | Mosquito | Japan | 1958 | LD50 | 4 | Yes |
| Notes: authors use units of LD50/0.4ml. | | | | | | | |
| Ilkal | P 20778 (human origin) | Mosquito | India | 1988 | LD50 | 7 | Yes |
| Johnsen | Mosquitoes in Thailand | Mosquito | Thailand | 1974 | PFU | 8 | No |
| Notes: the authors provided the viraemia for one pig and one dog as 100 PFU/ml and 3 PFU/ml, respectively. We assumed that the authors did not log transform this data before publication, so we took the base-10 logarithm of each value. | | | | | | | |
| Karna | Multiple, all from mozzies | Subcutaneous | US | 2019 | PFU | 30 | No |

| | | | | | | | |
|---|---|---|---|---|---|---|---|
| Notes: to account for the three ducks that did not develop a viraemia, we added a summarised group for Karna with three non-viraemic individuals into the dataset. | | | | | | | |
| van den Hurk | TS3306 (mosquito origin) | Mosquito | Australia | 2009 | TCID50 | 10 | No |
| Nemeth | JE-IN (human origin) | Subcutaneous | US | 2012 | PFU | 15 | No |
| Notes: individual viraemia data was not provided, but we felt that the information in Figure 2 was sufficiently detailed. From Figure 2 we extracted the highest titre value on the graph for each species and chose this value as the "mean peak viraemia" for all individuals of that species. Likewise, from Figure 2 we extracted the "average duration of viraemia" for each species by counting the number of days with viraemias above zero. Both extracted values may be overestimations, but we feel that the inclusion of this data is valuable. We also assumed that their ducks were domestic ducks (*Anas platyrhynchos domesticus*) given that they were sourced from a breeder. | | | | | | | |
| Cleton | JE-VN and JE-P3 (mosquito origin) | Subcutaneous | US | 2014 | PFU | 48 | No |
| Lunt | Nakayama (human origin) | Subcutaneous | Australia | 2001 | TCID50 | 6 | No |
| Park | JE-91 (mosquito origin) | Intravenously | US | 2018 | PFU | 5 | No |
| Ricklin | Nakayama (human origin) | Intravenously | Switzerland | 2016 | TCID50 | 12 | No |
| Ricklin | Nakayama (human origin) | Jugular vein, intradermal, and pig-to-pig | Switzerland | 2016 | Real-time RT–PCR, and expressed as U $ml^{-1}$ (1 U corresponding to the RNA quantity found in 1 TCID50 of a virus stock) | 5 | No |
| Scherer | "Pig 9" (pig origin); M5/596 (mosquito origin) | Subcutaneous | Japan | 1959 | LD50 | 4 | No |
| Xiao | 7 strains | Subcutaneous | China | 2018 | TCID50 | 10 | No |
